# Incidence of mortality and its predictors among adult Visceral Leishmaniasis patients at University of Gondar Comprehensive Specialized Hospital, Ethiopia

**DOI:** 10.1101/723379

**Authors:** Yigizie Yeshaw, Adino Tesfahun Tsegaye, Solomon Gedlu Nigatu

## Abstract

**Background:** Visceral leishmaniasis (VL) is a neglected tropical disease resulting in a huge burden of mortality and impact on development of a country. Even though anti-leishmanial drugs reduce the incidence of mortality among VL patients, still there is a death of VL patients while on treatment. However, study on incidence of mortality and its predictors among these patients while on treatment is scarce in Ethiopia.

**Objective:** The aim of this study was to determine incidence of mortality and its predictors among adult VL patients at University of Gondar Hospital.

**Methods:** Institution based retrospective follow up study was conducted from 2013 to 2018 at University of Gondar Hospital. Data were collected from patients’ chart and analyzed using Stata 14. Kaplan Meier failure curve and Log Rank test was used to compare survival probability of patients with categorical variables. Multivariable stratified Cox model was used to identify predictors of mortality among VL patients. P≤ 0.05 was employed to declare statistically significant factors. Adjusted Hazard Ratio (AHR) and its 95% confidence interval (95% CI) was estimated for potential risk factors included in the multivariable model.

**Results:** A total of 586 VL patients were included in the study. The median age of patients was 23 years. The incidence of mortality was 6.6 (95% CI: 5.2 - 8.4) per 1000 person-days of observation. Independent predictors of mortality were: presence of comorbidity (AHR=2.29 (95% CI: 1.27-4.11)), relapse VL (AHR=3.03 (95% CI: 1.25-7.35)), toxicity of treatment drug (AHR=5.87 (95% CI:3.30-10.44)), nasal bleeding (AHR=2.58 (95%CI: 1.48-4.51)), jaundice (AHR=2.84 (95% CI: 1.57-5.16)) and being bedridden (AHR=3.26 (95 % CI: 1.86-5.73)).

**Conclusion:** The incidence of mortality among VL patients was high. Mortality was higher among VL patients with concomitant disease, relapse, toxicity during treatment, nasal bleeding, jaundice, and bedridden patients. Therefore, strict follow up and treatment of VL patients who have comorbidity, relapse VL, toxicity, nasal bleeding and jaundice were crucial so as to reduce the risk of mortality.

**Authors’ summary:** Visceral leishmaniasis is a neglected tropical disease caused by a protozoa parasite. Over 90% of global burden of VL occurs in poor rural and suburban areas in seven countries including our country, Ethiopia. If not appropriately treated, over 95% of VL cases will eventually die. The emergence of VL in Ethiopia places a huge burden on society as it affects poor, young and productive age group of its population. However, there is scarcity of data about incidence of mortality and its predictors among adult VL patients in Ethiopia.

In this study, a registry of VL patients at Gondar University Hospital was taken to determine the incidence of VL mortality and its predictors. Mortality rate was higher among VL patients with concomitant disease, relapse, drug toxicity, nasal bleeding and jaundice. Therefore, strict follow up and treatment of VL patients who had comorbidity, relapse VL, drug toxicity, nasal bleeding and jaundice were crucial.

## Introduction

Visceral leishmaniasis (kal-azar) is a neglected tropical disease caused by a protozoa parasite called Leishmania donovani complex (L. donovani and L. infantum), which is transmitted by the female phlebotomine sand flies. It is characterized by prolonged fever, weight loss, decreased appetite, anemia, and hepatosplenomegaly[1–3].

Globally about 500,000 new cases of visceral leishmaniasis (VL) occur every year. Of these, over 90% of global burden of VL occurs in poor rural and suburban areas in seven countries: Bangladesh, Brazil, Ethiopia, India, Nepal, Sudan and South Sudan[2,4].

From Eastern Africa region, Sudan is the most affected country, followed by Ethiopia, Kenya, Somalia and Uganda[5]. According to a study in eastern Uganda, mortality rate among VL patients is 3.7% [6].

When we came to Ethiopia, mortality rate among severely ill adult VL patients is 4.8 % [7] and it is 14 % among VL-HIV co-infected adults [8]. Another two cross sectional studies shoes that the proportion of death among VL patients is 12.4% in Kahsay Abera Hospital [9] and 18.5% in Tigray region [10].

Visceral leishmaniasis is associated with about 2,357,000 disability-adjusted life years (DALYs), placing leishmaniasis ninth in a global analysis of infectious diseases [3,11]. If not appropriately treated, over 95% of VL cases will eventually die, resulting in at least 50,000 deaths per year worldwide, a rate surpassed among parasitic diseases only by malaria [12]. In recent years, more effective treatments have reduced the case fatality rate to 10% on average, which is equivalent to a fatality rate of 20,000 – 40,000 deaths[13].

The emergence of VL in Ethiopia places a huge burden on society in terms of mortality, and impact on country’s economy as well as future development. This is because the disease is more prevalent in Kola to Weina Dega agro-ecological zones of Ethiopia, areas of major agricultural projects such as dams for electricity and irrigation purposes as well as other agricultural activities all exist [14].

Predictors of mortality among VL patients include: presence of drug toxicity [15], malnutrition [3,16,17], VL-HIV co-infection [5,9,17–24],thrombocytopenia [5,11,18–21,25], leukopenia [5,18–21,26], jaundice [5,18–21,26], relapsing course of the disease [11,22,25], high parasite load [8,27,28], renal failure(creatinine>1.5 mg/dl)[20,26], diarrhea [10,11,25], bleeding [5,10,18–21], anemia [11,17,25], inability to walk at admission [9], longer duration of illness [17], concomitant disease [5,18–21], late diagnosis [6,9,29] and edema [5,18–21].

Even though visceral leishmaniasis has become one of the leading health problem in Ethiopia as it causes high mortality rate and reduced productivity by affecting a significant portion of poor, rural, and productive age group of the country [30], there is scarcity of data about incidence of mortality and its predictors among adult VL patients. Hence, considering VL severity and lethality rate as well as its impact on a country, early identification of factors associated with mortality among VL patients is relevant to the premature establishment of appropriate measures. Therefore, the objective of this study is to determine incidence of mortality and its predictors among adult VL patients who were on treatment at University of Gondar Comprehensive Specialized Hospital from 2013 to 2018.

## Methods

### Study area and period

This study was conducted at University of Gondar Comprehensive Specialized Hospital, Kal-azar ward from January 1, 2013 to December 30, 2018.

University of Gondar Hospital is located 727 km far from the capital city, Addis Ababa in northwest direction. It serves for a population of five million across the region.

The Hospital has Leishmaniasis Research and Treatment Center (LRTC), which was established in 2004 in collaboration with Drugs for Neglected Diseases initiatives (DNDi). In this center, several clinical trials have been conducted. The staff at LRTC are trained and experienced in good clinical practice to treat all forms of leishmaniasis. Visceral leishmaniasis suspected patients are referred from the different units of the Hospital and from other health facilities of the catchment area. Patients admitted for VL treatment in LRTC are routinely evaluated and the findings are documented in their chart records. The LRTC currently serves for more than 300 VL patients per year[7].

### Study design

Institution based retrospective follow up study was employed.

### Source and study population

All adult VL patients who were treated with anti-leishmanial drugs at University of Gondar Hospital were the source population and all adult VL patients who were treated with anti-leishmaniasis drug from 2013 to 2018 were the study population.

### Inclusion and exclusion criteria

All adult VL patients who were treated with anti-leishmaniasis drug from Jan 2013 to Dec 2018. Patients with unknown treatment outcome, no recorded date of treatment initiation and treatment outcome were excluded.

### Sample size and sampling procedure

The sample size for this study was calculated through Stata 14 software using 12.4 % probability of an event (death) in another similar setting [9], 80 % power, hazard ratio of two, 5% significance level, and 10 % for incomplete data. Accordingly, the final sample size of the study was 586. Simple random sampling technique was used to select these sample patients’ charts from total of 1899 patients that had been on treatment from 2013 to 2018.

### Study variables

The dependent variable for this study was time until death of the patient. Independent variables include socio-demographic variables (age, sex, residence, migration status), clinical and laboratory related variables such as visceral leishmaniasis parasite load, leukopenia, thrombocytopenia, hemoglobin level, treatment type, toxicity during treatment, late diagnosis, VL episode, concomitant disease, condition of patient at admission, creatinine level, diarrhea, jaundice, BMI, nasal bleeding and edema.

### Operational definitions

#### Event (Death)

any documented death of VL patient while taking the treatment during follow up period.

#### Censored

Patients who were transferred out, treatment failure or loss to follow up or became initial cured.

#### Initial cure

declared when a patient shows an improvement of signs and symptoms at the end of treatment depending on the category of treatment (after 12 days for those patients taking ambisome, 17 days for those patients taking a combination of SSG and PM, and 28 days for those patients taking SSG only) such as fever resolution, hemoglobin increase, weight gain and spleen size regression), and/or has a negative parasitological test of cure (TOC).

#### Treatment failure

defined as a positive TOC (parasitological failure) and/or persisting clinical signs/symptoms at the end of treatment or failure to continue first-line treatment for safety reasons.

#### Loss to follow up

A patient who started VL treatment but interrupted treatment due to the patient leaving the Hospital during the study period [9,15,31].

#### Primary VL case

a patient who is diagnosed with visceral Leishmaniasis for the first time in which diagnosis relies on a positive serological test for VL (rK39 based rapid test and/or DAT direct agglutination test) and/or a positive parasitological test (microscopic detection of Leishmania parasites in splenic aspirate).

#### Relapse VL

a patient with a history of previous VL and discharged improved or with a negative test of cure (TOC) after treatment and who then presents with symptoms of VL after four weeks of initial VL treatment and is parasitologically confirmed and documented as relapse VL[32].

**High parasite load** is parasite load grade of more than 4+ (1–10 parasites per field). Low parasite load is parasite load grade of less than or equal to 3(1–10 parasites per 10-1000 fields).

#### Concomitant disease

presence of one or more of a documented case of the following diseases: tuberculosis, pneumonia, malaria and HIV.

#### Toxicity during treatment

presence of one or more of documented toxicity such as cardiac arrest, pancreatitis, jaundice (liver disease) and kidney failure [20].

### Data collection procedure and tools

Data were collected from VL patient charts who were registered from 2013 to 2018 using pretested and structured data extraction checklist.

Based on the pretest finding, amendments and arrangements were made on the data extraction checklist. Four BSc Nurse data collectors were recruited and trained for half day about ways of extracting data from patient charts. Clinical and laboratory parameters: such as parasite load, leukopenia, thrombocytopenia, hemoglobin level, treatment type, late diagnosis, VL episode, concomitant disease, general condition of patient at admission, diarrhea, jaundice, BMI, nasal bleeding, edema, and creatinine level were extracted at admission. The presence of toxicity during treatment was also assessed. The presence or absence those abnormalities was decided based on the documentation made by the physicians. Laboratory results were also collected at admission and their value was compared with their reference value to decide on the presence of derangement on these parameters. Patients were followed retrospectively for 12 to 28 days according to their treatment category.

### Data quality management

To assure the data quality, high emphasis was given in designing data collection instrument. Training was given for data collectors to create a common understanding of the data extraction checklist and patient chart as well as registry reviewing skills. The data extraction checklist was pre-tested. Throughout the course of the data collection, data collectors were supervised by the principal investigator.

### Data analysis procedure

Data were checked on a daily basis for completeness, clarity and accuracy. Data were entered into Epi-data version 3.1 and exported to Stata 14 for analysis. Descriptive measures such as frequency of each categorical variable was calculated.

Time to event data (survival times in days) was calculated by subtracting date of treatment (t_o_) started from date of event occurred (t_1_). Death was the outcome variable (event) that was measured (coded as 1 for death and 0 for censored). Person-days of observation (PDO) were calculated from the date of starting anti-leishmanial treatment to date of death or censored. The failure probability of patients during VL treatment with respect to socio-demographic and clinical variables was described with the Kaplan Meier (KM) curve. Log-rank test was also used to test the failure differences among categories of each independent variable.

Schoenfeld residuals test (both global and scaled) and graphical methods were used to check the Cox proportional hazard assumption. Model adequacy was also checked using Cox Snell residuals.

Multivariable stratified Cox model was used to identify predictors of mortality among VL patients. All variables with a p-value of < 0.2 at bi-variable analysis were entered into the final model. P-Value ≤ 0.05 was employed to declare the statistically significant variables. Adjusted hazard ratio (AHR) and its corresponding 95% confidence interval (95% CI) were estimated for potential risk factors included in the multivariable stratified Cox model.

### Ethical consideration

Ethical clearance was obtained from Institutional Review Board of Institute of Public Health, University of Gondar (Reference number IPH/180/06/2011). Written permission letter for reviewing extracting data from patient’s chart was also obtained from University of Gondar Comprehensive Specialized Hospital. Privacy and confidentiality of information was kept properly and names of patients as well as other personal identifiers were not recorded.

## Results

### Sociodemographic and baseline clinical characteristics of VL patients

A total of 586 visceral leishmaniasis patients were included in the study. Almost all, 584(99.7%) of them were males. The median age of patients was 23 years with interquartile range of 20 to 28 years. Most of the patients, 470 (80.2%) were migrant workers.

Majority of VL patients, 561(95.7%) had primary visceral leishmaniasis. From a total of 586 patients, 169 (28.8 %) of them had concomitant disease at admission. Of these, about half of them had pneumonia (49.7%). Sixty-seven (11.4 %) of them had high parasite load. Forty-one (7.0%) of study participants had toxicity during treatment. Of these, 15(36.6 %) of them had cardiac arrest followed by pancreatitis, 10(24.4 %). Regarding duration of illness, 258(44%) of them had more than 30 days of illness duration at admission **(Table 1)**.

**Table 1:**
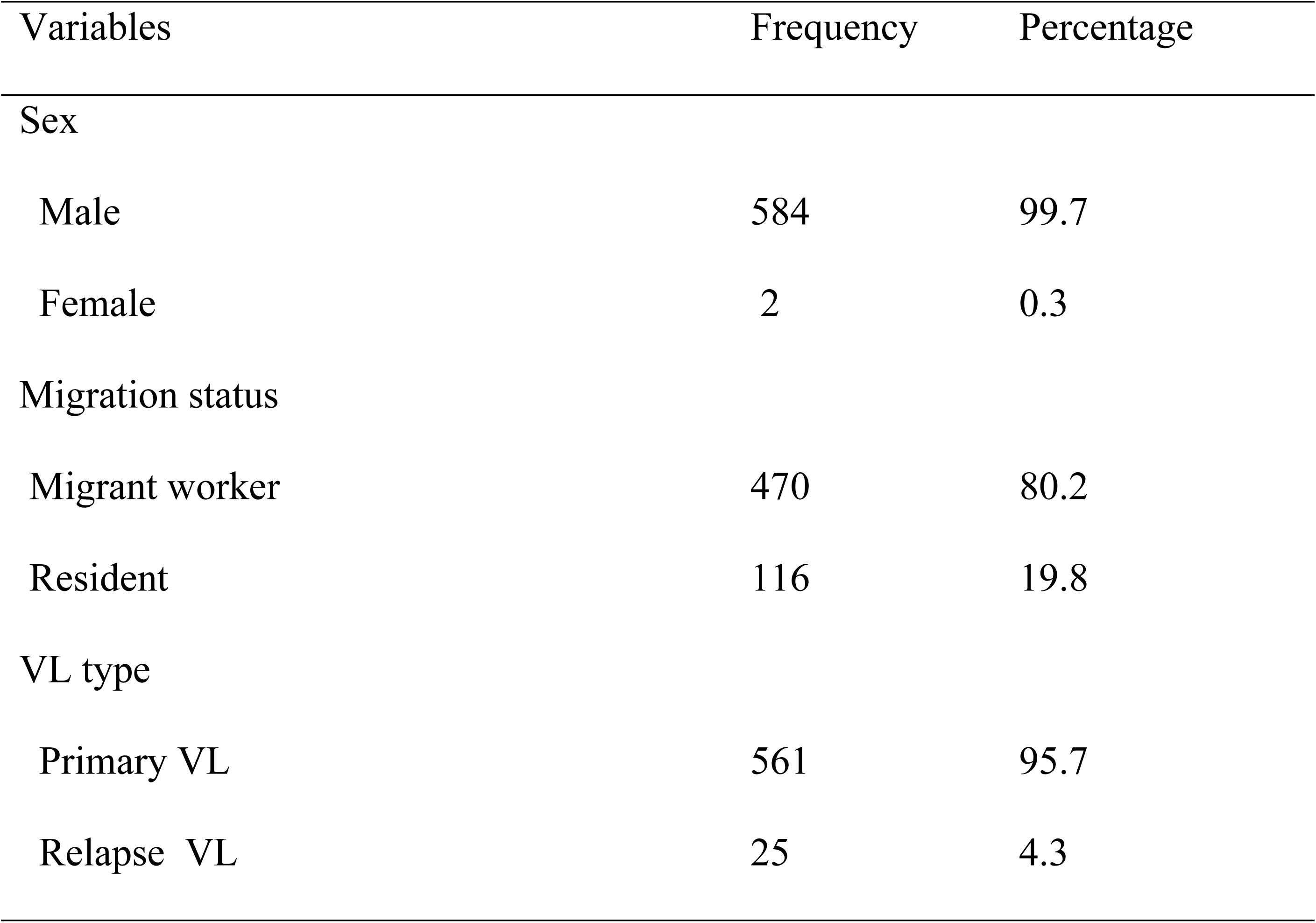
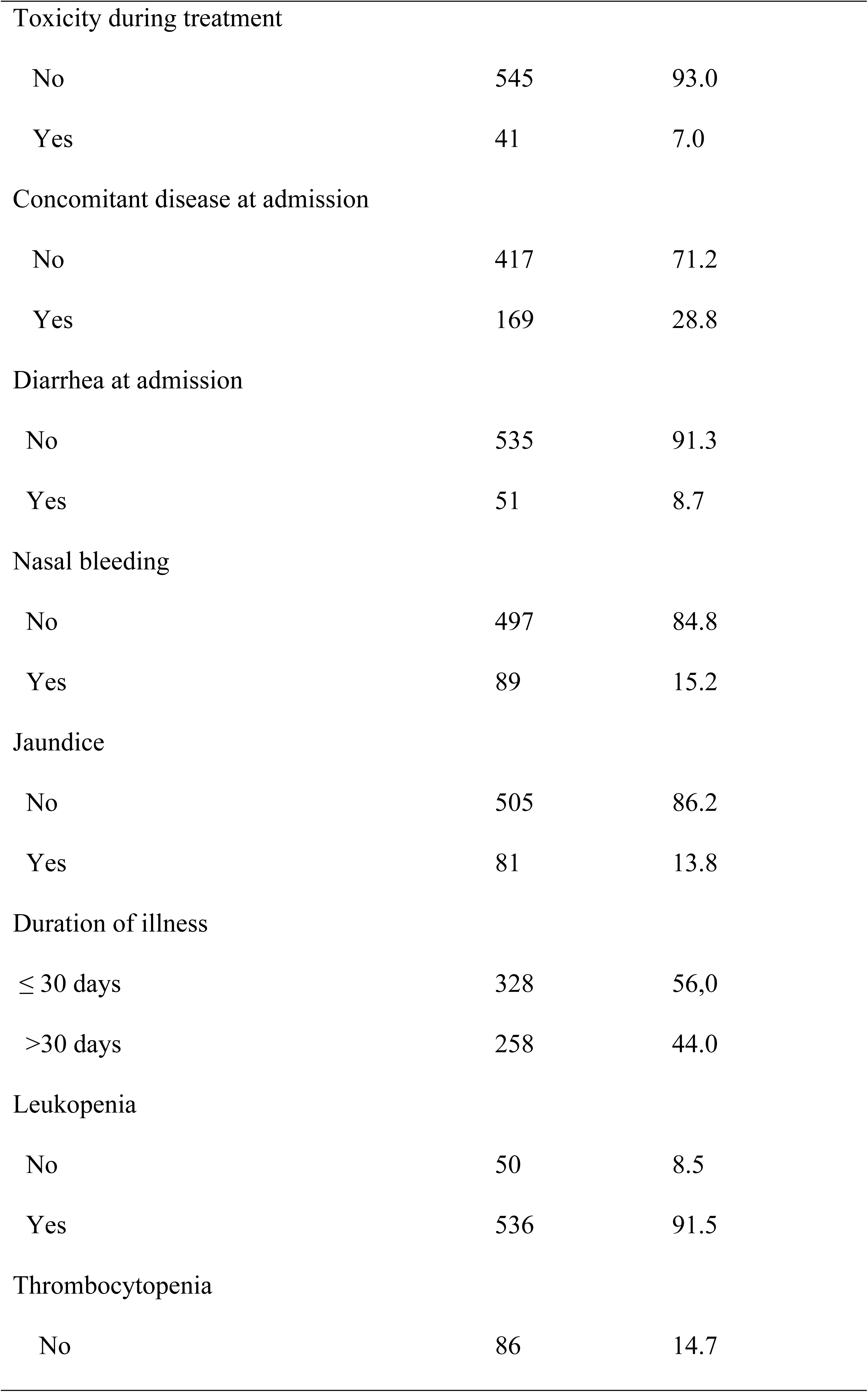
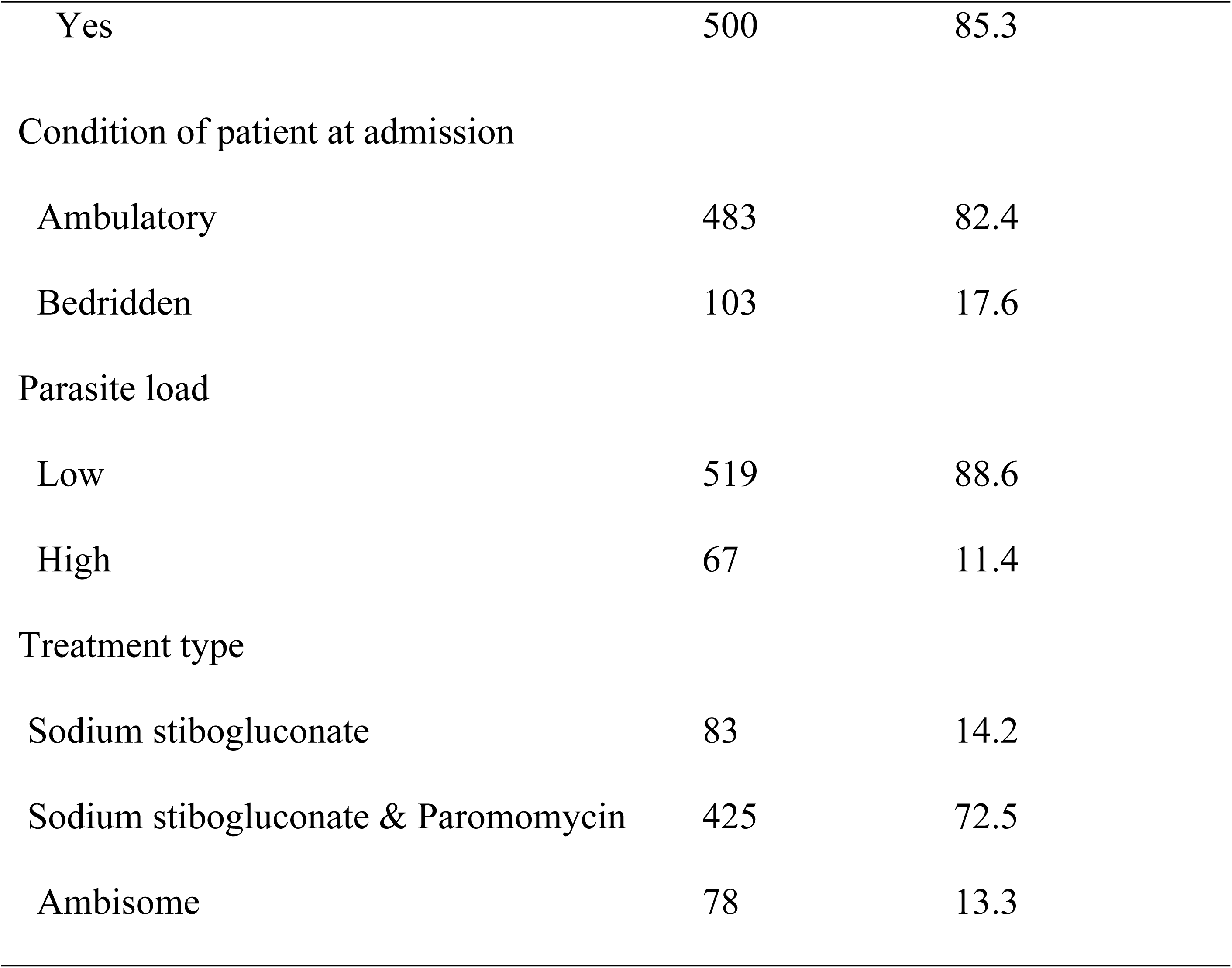
socio-demographic and baseline clinical characteristics of VL patients at University of Gondar Hospital, 2019(n=586)

### Incidence of mortality among VL patients

From the total of 586 VL patients who start anti-leishimanial treatment during the study period, 65 (11.09 %) of them were died, 483(82.4 %) cured, 26(4.4%) lost to follow up, 9(1.5%) treatment failure, and the rest, 3 (0.5%) were transferred out. The total cohort contributed 9830 person-days, resulting in the overall mortality rate of 6.6 deaths (95% CI: 5.2 - 8.4) per 1000 person-days of observation. Of the 65 deaths, 39 (60%) of them occurred within the first 10 days of treatment initiation, which is equivalent to a mortality rate of 6.9 deaths per 1000 person-days of observation. The cumulative failure probability of VL patients at the end of follow up period was 0.193 **(Figure 1)**.

**Figure 1:**
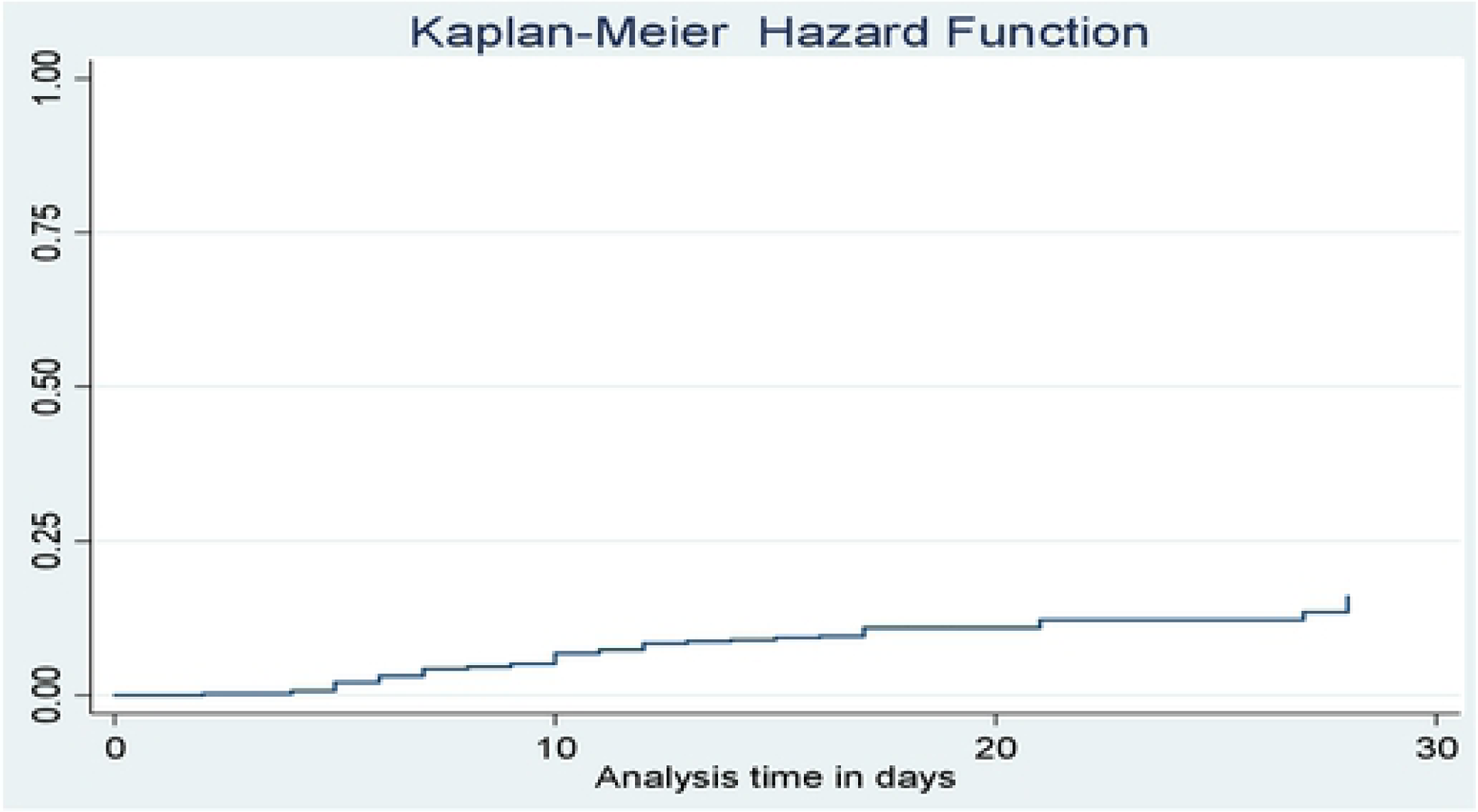
Cumulative hazard function of VL patients at UoG Hospital, 2019.

### Comparison of failure functions

Kaplan Meier failure curve was used to compare death probability among categories of each independent variable visually. Log-rank test was also used to objectively judge the presence or absence of a difference in death probabilities among different categories of each independent variable. Accordingly, Kaplan Meier failure curve was done for all possible predictors. For instance, relapse VL patients had shorter survival experience than primary VL cases. This visually observed difference was also statistically significant (Log rank, p<0.001). Visceral leishmaniasis patients who had comorbidity at admission had shorter survival experience than those VL patients without comorbidity (Log rank, p<0.001). VL patient who had jaundice at admission had shorter survival experience than those VL patients without jaundice (Log rank, p<0.001) (**Figure 2**).

**Figure 2:**
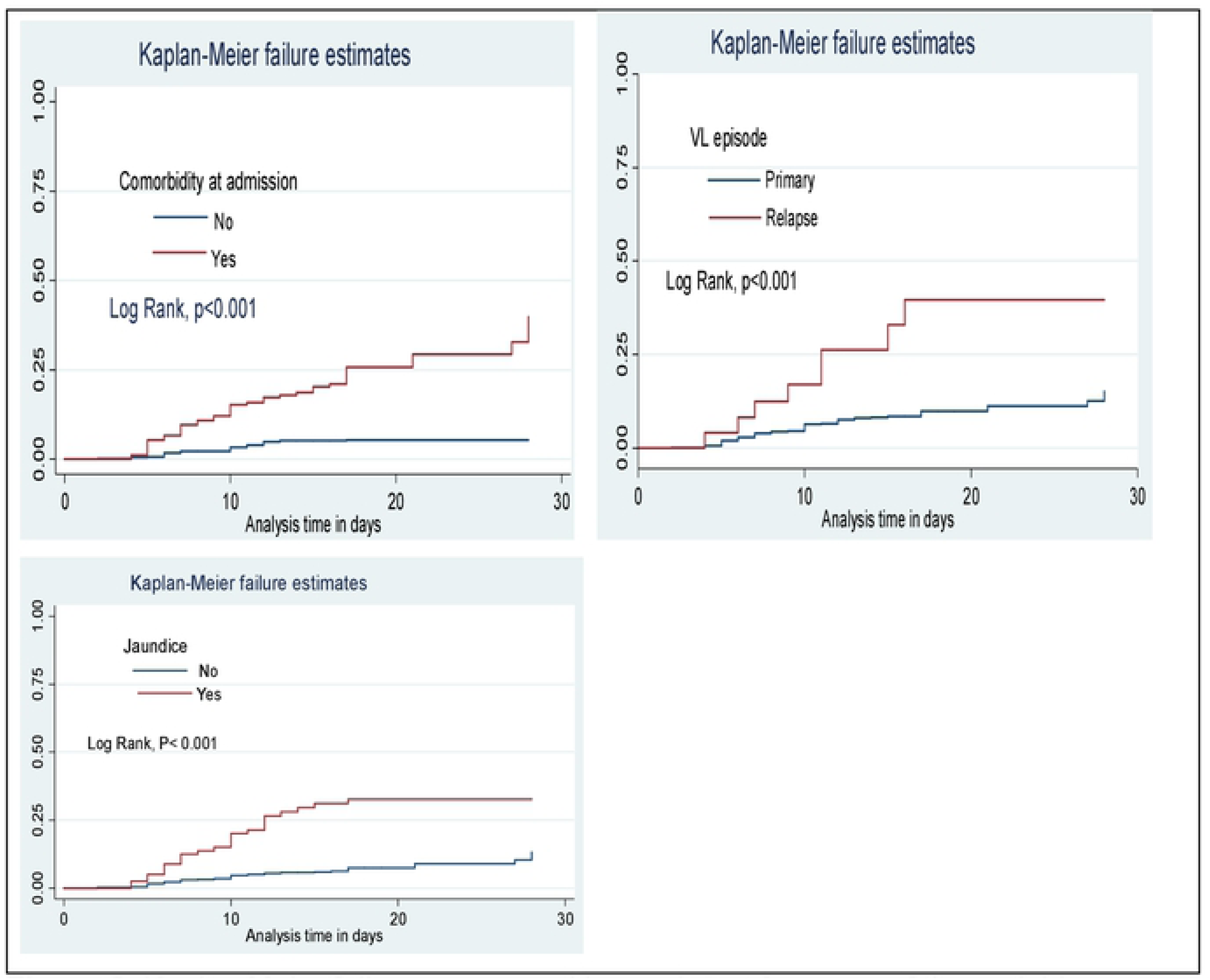
Kaplan Meier failure curves and log rank test for some of the variables among the cohort of VL patients at UoG Hospital, 2019.

### Assessing Proportional Hazard Assumption

Proportional hazard assumption was checked both graphically and Schoenfeld residuals test (global and scaled) for all possible predictors of mortality. Just to show for some of the variables –Ln (-Ln (survival probability) to Ln (analysis time) for jaundice, comorbidity and residence was demonstrated graphically. Accordingly, the hazards do not cross between categories of jaundice and comorbidity, which means that the proportional hazard assumption was satisfied for these variables. However, it crosses between categories of residence, which means that proportional hazard assumption was not satisfied for residence (**Figure 3**).

**Figure 3:**
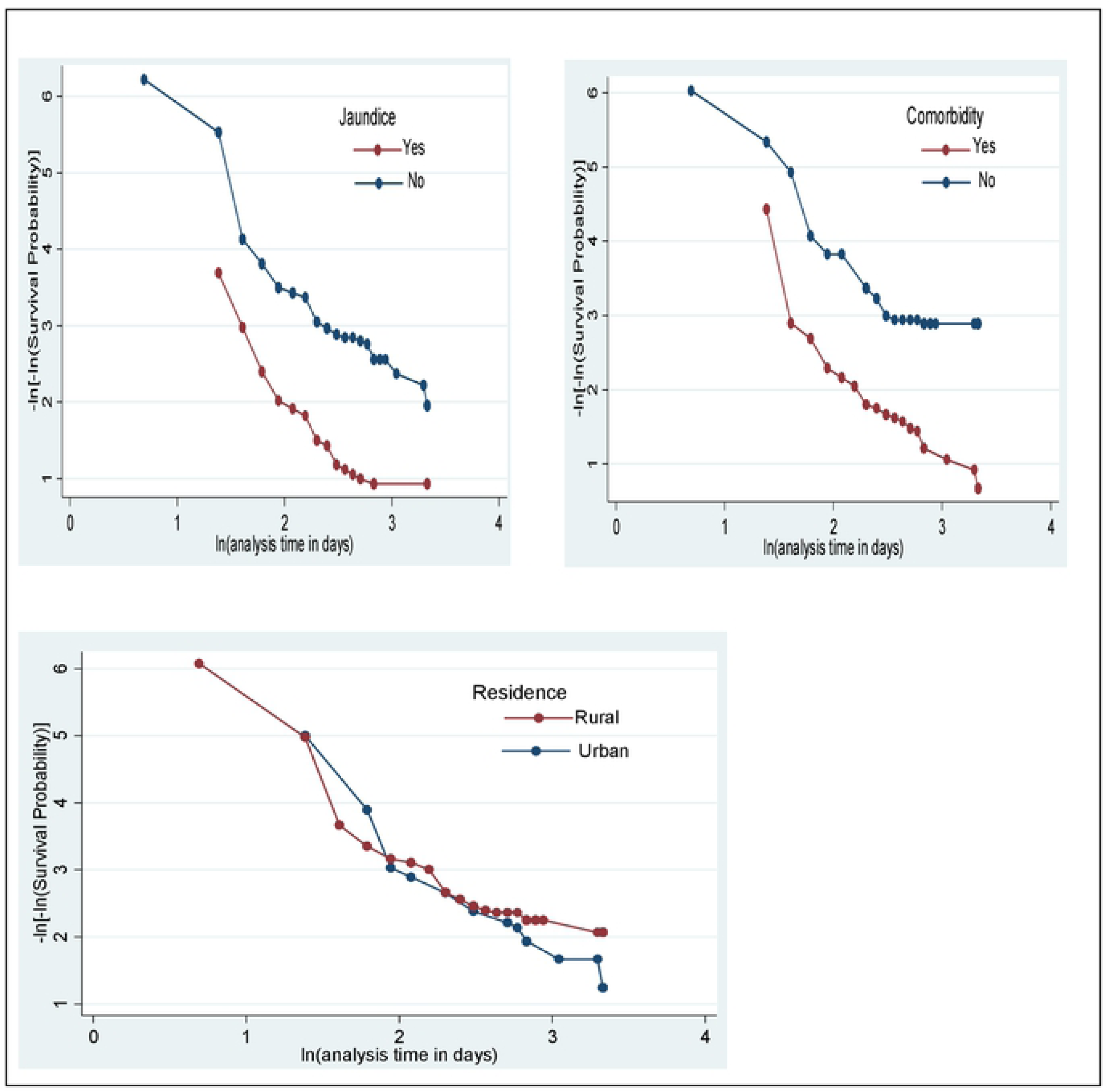
Proportional hazard plot for some of the variables among the cohort of VL patients at UoG Hospital, 2019.

Moreover, in order to test proportional hazard assumption objectively, Schoenfeld residuals test (global and scaled) was done. Accordingly, all variables with the exception of residence satisfies proportional hazard assumption (p>0.05) (**Table 2**).

**Table 2:** Proportional hazard assumption test for the study on incidence of mortality and its determinants among VL patients at UoG Hospital, 2019

| Variables | rho | Chi2 | df | Prob>chi2 |
| --- | --- | --- | --- | --- |
| Residence | -0.29213 | 6,20 | 1 | 0.0127 |
| Edema | -0.19109 | 2.73 | 1 | 0.0983 |
| Diarrhea | 0.06225 | 0.28 | 1 | 0.5940 |
| Comorbidity | 0.16637 | 2.29 | 1 | 0.1305 |
| VL episode | -0.15490 | 1.77 | 1 | 0.1831 |
| Toxicity | -0.01293 | 0.01 | 1 | 0.9055 |
| Hemoglobin | -0.10120 | 0.78 | 1 | 0.3781 |
| Nasal bleeding | 0.06777 | 0.38 | 1 | 0.5398 |
| Jaundice | -0.05067 | 0.19 | 1 | 0.6626 |
| Illness duration | 0.14165 | 1.61 | 1 | 0.2039 |
| Patient condition at admission | -0.11546 | 1.22 | 1 | 0.2688 |
| Parasite level | 0.13763 | 1.63 | 1 | 0.2011 |
| Creatinine level | 0.02843 | 0.06 | 1 | 0.8060 |
| Treatment type | 0.04827 | 0.17 | 1 | 0.6793 |
| Global test |  | 15.30 | 14 | 0.4299 |

### Determinants of mortality among VL patients

Variables with p<0.2 on bivariable analysis were entered to multivariable stratified Cox model and six variables were found to be an independent predictor’s of mortality among VL patients while on treatment (p≤ 0.05). These include: concomitant disease, episode of visceral leishmaniasis, toxicity during treatment, nasal bleeding, jaundice, and patient condition at admission.

The hazard of death among relapse VL patients was 3 (AHR=3.03(95%CI: 1.25-7.35)) times higher than primary VL patients. The risk of death was 5.9 (AHR=5.87(95%CI: 3.30-10.44)) times higher among patients who had toxicity during treatment as compared to those patients who didn’t have toxicity. The hazard of death among VL patients with comorbidity was 2.3 (AHR=2.29(95% CI: 1.27-4.11)) times higher than those who didn’t have. The hazard of death among VL patients who had nasal bleeding was 2.6 (AHR=2.58(95%CI: 1.48-4.51)) times higher than those patients who didn’t have nasal bleeding. Visceral leishmaniasis patients who had jaundice at admission were 2.8 (AHR=2.84(95%CI: 1.57-5.16)) times more at risk of death than their counterparts. Those patients who were bedridden had 3.3 (AHR=3.26(95%CI: 1.86-5.73)) times increased risk of death compared to ambulatory patients (**Table 3**).

**Table 3:** Multivariable stratified Cox regression analysis for incidence of mortality among VL patients at UoG Hospital Gondar, 2019.

| Variables | Death |  | Hazard Ratio (95 % CI) |  |
| --- | --- | --- | --- | --- |
|  | No | Yes | CHR | AHR |
|  | N (%) | N (%) |  |  |
| Age |  |  | 1.04(1.01-1.07) | 0.99 (0.95- 1.04) |
| VL type |  |  |  |  |
| Primary VL | 504 (89.8) | 57(10.2) | 1 | 1 |
| Relapse VL | 17(68) | 8(32) | 3.98(1.89- 8.36) | 3.03(1.25-7.35)* |
| Hemoglobin |  |  |  |  |
| 0-7.9 mg/dl | 218(85.5) | 37(14.5) | 1.59(0.71- 3.57) | 1.42(0.59-3.42) |
| 8-10.9 mg/dl | 232(91.7) | 21(8.3) | 0.87(0.37-2.05) | 1.06(0.42-2.56) |
| ≥11 mg/dl | 71(91.0) | 7(8.9) | 1 | 1 |
| Illness duration |  |  |  |  |
| ≤30 days | 300(91.5) | 28(8.5) | 1 | 1 |
| >30 days | 221(85.7) | 37(14.3) | 1.64(1.01-2.69) | 0.92(0.51-1.64) |
| Rx toxicity |  |  |  |  |
| No | 507(93) | 38(7) | 1 | 1 |
| Yes | 14(34.1) | 27 (65.9) | 13.99(8.48-23.10) | 5.87(3.3-10.44)* |
| Comorbidity |  |  |  |  |
| No | 395(94.7) | 22(5.3) | 1 | 1 |
| Yes | 126(74.6) | 43(25.4) | 5.57(3.33- 9.32) | 2.29(1.27-4.11)* |
| Diarrhea |  |  |  |  |
| No | 482(90.1) | 53(9.9) | 1 | 1 |
| Yes | 39(76.5) | 12(23.5) | 2.61(1.39- 4.89) | 1.73(0.88-3.78) |
| Nasal bleeding |  |  |  |  |
| No | 459(92.3) | 38(7.7) | 1 | 1 |
| Yes | 62(69.7) | 27(30.3) | 4.51(2.75- 7.39) | 2.58(1.48-4.51)* |
| Edema |  |  |  |  |
| No | 443(90.8) | 45(9.2) | 1 | 1 |
| Yes | 78(79.6) | 20(20.4) | 2.34(1.38 -3.97) | 1.45(0.78-2.66) |
| Jaundice |  |  |  |  |
| No | 465 (92.0) | 40(8.0) | 1 | 1 |
| Yes | 56(69.1) | 25( 30.9) | 4.62(2.79- 7.62) | 2.84(1.57-5.16)* |
| Creatinine level |  |  |  |  |
| <1.5 mg/dl | 455(91.0) | 45(9.0) | 1 | 1 |
| ≥1.5mg/dl | 66(76.7) | 20(27.3) | 2.88(1.70- 4.88) | 1.33(0.70-2.54) |
| Condition of patient |  |  |  |  |
| Ambulatory | 456(94.4) | 27(5.6) | 1 | 1 |
| Bedridden | 65(63.1) | 38(36.9) | 8.29(5.05- 13.59) | 3.26(1.86-5.73)* |
| Parasite load |  |  |  |  |
| Low | 469(90.4) | 50(9.6) | 1 | 1 |
| High | 52 (77.6) | 15(22.4) | 2.29(1.28- 4.10) | 1.94(0.97-3.89) |
| Treatment type |  |  |  |  |
| SSG and PM | 398(93.7) | 27(6.3) | 1 | 1 |
| SSG | 69(83.1) | 14(16.9) | 2.16(1.07- 4.34) | 0.81(0.37-1.78) |
| Ambisome | 54(69.2) | 24(30.8) | 7.82(4.39- 13.89) | 1.79(0.91-3.54) |

SSG: Sodium stibogluconate, PM: Paromomycin, \*p≤0.05

## Discussion

This study aimed to identify incidence of mortality and its predictors among adult VL patients on treatment. Accordingly, the overall incidence rate of mortality was 6.6 (95% CI 5.2 - 8.4) per 1000 person-days of observation, with most of deaths (60%) occurring within the first 10 days of follow-up period, which requires the attention of health workers in these periods. The proportion of death among VL patients in this study was 11.09 % (95% CI: 10.85% - 13.6%), which is in line with a study conducted in Kahsay Abera Hospital(12.4%) [9]. This similarity might be due to the similarity in quality of care given for VL patients in these Hospitals, as these Hospitals use similar visceral leishmaniasis treatment guideline. Moreover, most of the patients in the current and earlier study were migrant workers as well as living in a rural area, which share similar economic level.

However, this finding is less than the finding in Tigray(18.5%)[10] as well as a cross-sectional study conducted among VL-HIV co-infected adults of Ethiopia(14.0%) [8]. The possible explanation for this difference might be differences in type of antileishmanial drugs used, in which patients in the earlier studies used only Sodium stibogluconate, which is often poorly tolerated, and toxic drug, to result in a significant incidence of serious adverse events such as toxicity of pancreas, liver, kidney and heart than other anti-leishimanial drugs [33]. Moreover, presence of comorbidity (VL-HIV co-infection), may increase the risk of death in the previous study compared to the current one.

In the contrary, the current finding is higher than the finding of Eastern Uganda (3.7%)[6], and Northwest Ethiopia (4.8 %) [7]. The reason for this discrepancy in the case of Eastern Uganda might be that only primary VL cases were included in the study, which may underestimate death rate, as death is more common among relapse cases than primary VL patients[22,25]. In the case of earlier study of Ethiopia, VL patients who were taking Amphotericin B only were included, a drug with less toxicity and more tolerability than that took by participants of this study such as Sodium stibogluconate [12].

In the current study, the hazard of death among relapse VL patients was 3 times higher than that of primary VL patients. This finding is similar with the finding of two studies in Brazil [22,25]. This might be because of majority of the relapse cases in this study were HIV patients (64%). Since both VL and HIV attack the immune system of the body, they produce a profound immune deficiency state. The results and effect of this state is that VL speeds up the onset of full-blown AIDS and shortens the life expectancy of HIV-infected individuals, while HIV complicates management of VL. Visceral leishmaniasis lowers the total lymphocyte count (TLC) and Cluster of Differentiation four(CD4) count to a great extent by depressing the bone marrow and the splenic activities[34].

In our study, VL patients who had toxicity during treatment were 5.8 times more at risk of mortality than those patients who had no toxicity. This finding is similar with the finding of studies in Uganda, Somali and Ethiopia [6,11,12]. Since most anti-leishmanial drugs are toxic, development of drug toxicity such as arrhythmia, pancreatitis and others is common, which lead to poor compliance and further deterioration of the patient to cause death[15].

The hazard of death among VL patients with comorbidity was 2.3 times higher than those without comorbidity. This finding is in agreement with the finding of studies in Eastern Uganda, Brazil, India, and Ethiopia [6,22–24,29]. This might be due to the double burden associated with the comorbidity. Moreover, patients with concomitant disease/ comorbidity at admission had to take more drugs so they might have more risk of toxicity and drug-drug interaction, which causes severe form of the disease to end up with death of the patient.

The hazard of death among VL patients who had nasal bleeding was 2.6 times higher than those patients who didn’t have nasal bleeding. This result is consistent with the result of a study in Northern Ethiopia(Tigray), America and Sudan[5,10,21]. Nasal bleeding among VL patients occurs probably due to a combination of deficient clotting factor and platelet count, which increases the risk of death among VL patients[35].

In this study, VL patients who had jaundice at admission were 2.9 times more at risk of death than their counterparts. The finding of this study is similar with studies conducted in Gedaref state of Sudan, America and Brazil [5,18–21,26]. This could be possibly due to, jaundice (usually the sign of liver dysfunction) causes decreased plasma protein synthesis, inability to detoxify drugs and impairment of other liver functions, to cause an increased risk of death. Those patients who were bedridden had 3.2 times increased risk of death compared to ambulatory patients. The current finding is similar with the finding of a study in Kahsay Abera Hospital [9]. This might be due to majority of bedridden patients (64%) in this study had concomitant diseases such as HIV/AIDS, tuberculosis and pneumonia, which ultimately increases the risk of death compared to ambulatory patients. This explanation is supported by studies conducted in Brazil, which states that most severely ill patients have an increased risk of concomitant diseases that can increase their risk of death [18,36]. Furthermore, severely ill patients usually do not respond to their medication easily and do not take adequate food as well. As limitation, since the study is retrospective, data about some variables such as blood glucose level, serum albumin level and income of the patient was not collected. So we can’t assess the effect of these variables on the incidence of mortality.

## Conclusion

The incidence of mortality among VL patients on treatment was high. The risk of death were higher among VL patients with concomitant disease, relapse, treatment toxicity, nasal bleeding, jaundice and those who were bedridden at admission, which requires greater attention for those risk groups.

## Recommendations

Health professionals and Hospital managers better to strictly follow and treat VL patients who had toxicity during treatment, nasal bleeding, jaundice, relapse and bedridden. They also give high emphasis for VL patients with other comorbidities such as pneumonia, HIV/AIDS, and tuberculosis. Conducting prospective cohort study by including the above missed variables is also recommended by the next researchers.

## Acknowledgments

I want to thank University of Gondar for giving this chance to conduct the study. My honest gratitude also goes to my friends for their valuable support and help. I want to thank my data collectors and chart room workers.

## Author Contributions

Conceptualization: Yigizie Yeshaw, Adino Tesfahun, Solomon Gedlu

Data cu ration: Yigizie Yeshaw, Adino Tesfahun, Solomon Gedlu

Formal analysis: Yigizie Yeshaw

Investigation: Yigizie Yeshaw, Adino Tesfahun, Solomon Gedlu

Methodology: Yigizie Yeshaw, Adino Tesfahun, Solomon Gedlu

Resources: Yigizie Yeshaw

Validation: Yigizie Yeshaw, Solomon Gedlu, Adino Tesfahun

Writing – original draft: Yigizie Yeshaw

Writing – review & editing: Solomon Gedlu, Adino Tesfahun

## Abbreviations/Acronyms

AHR: Adjusted Hazard Ratio
AIDS: Acquired Immune Deficiency Syndrome BMI Body Mass Index
CBC: Complete Blood Count
HIV: Human Immunodeficiency Virus
LRTC: Leishmaniasis Research and Treatment Center PM Paromomycin
RBC: Red Blood Cell
SSG: Sodium stibogluconate
TOC: Test of Cure
UoG: University of Gondar
VL: Visceral Leishmaniasis
WBC: White Blood Cell

## References

1. Biologics I for international cooperation in animal. Leishmaniasis Leishmaniasis (cutaneous and visceral). Cent food Secur public Heal. 2017;1–18.

2. WHO FR. Book Review: Working to Overcome the Global Impact of Neglected Tropical Diseases. Perspect Public Health. 2012;132(4):192–192.

3. Marinkelle CJ. The control of leishmaniases. Bull World Health Organ. 2010;58(6):807–18.

4. Alvar J, Vélez ID, Bern C, Herrero M, Desjeux P, Cano J, et al. Leishmaniasis worldwide and global estimates of its incidence. PLoS One. 2012;7(5):1–12.

5. Mueller YK, Nackers F, Ahmed KA, Boelaert M, Djoumessi JC, Eltigani R, et al. Burden of Visceral Leishmaniasis in Villages of Eastern Gedaref State, Sudan: An Exhaustive Cross-Sectional Survey. PLoS Negl Trop Dis. 2012;6(11):1–6.

6. Mueller Y, Mbulamberi DB, Odermatt P, Hoffmann A, Loutan L, Chappuis F. Risk factors for in-hospital mortality of visceral leishmaniasis patients in eastern Uganda. Trop Med Int Heal. 2009;14(8):910–7.

7. Tamiru A, Tigabu B, Yifru S, Diro E, Hailu A. Safety and efficacy of liposomal amphotericin B for treatment of complicated visceral leishmaniasis in patients without HIV, North-West Ethiopia. BMC Infect Dis [Internet]. 2016;16(1):1–7. Available from: http://dx.doi.org/10.1186/s12879-016-1746-1

8. Diro E, Lynen L, Mohammed R, Boelaert M, Hailu A, van Griensven J. High Parasitological Failure Rate of Visceral Leishmaniasis to Sodium Stibogluconate among HIV Co-infected Adults in Ethiopia. PLoS Negl Trop Dis. 2014;8(5):14–7.

9. Welay GM, Alene KA, Dachew BA. Visceral leishmaniasis treatment outcome and its determinants in northwest Ethiopia. Epidemiol Health [Internet]. 2016;39:1–6. Available from: http://www.e-epih.org/journal/view.php?doi=10.4178/epih.e2017001

10. Lyons S, Veeken H, Long J. Visceral leishmaniasis and HIV in Tigray, Ethiopia. Trop Med Int Heal. 2003;8(8):733–9.

11. Director H promotion and D prevention DG. Guidline for diagnosis, treatment and prevention of leishmaniasis in Ethiopia. 2017;91:1–88.

12. Government SF. Guidelines for diagnosis, treatment and prevention of visceral leishmaniasis in Somalia 2012. 2012;4–82.

13. Alvar J, Vélez ID, Bern C, Herrero M, Desjeux P, Cano J, et al. Leishmaniasis worldwide and global estimates of its incidence. PLoS One. 2012;7(5):1–12.

14. Gadisa E, Tsegaw T, Abera A, Elnaiem DE, Den Boer M, Aseffa A, et al. Eco-epidemiology of visceral leishmaniasis in Ethiopia. BMC [Internet]. 2015;8(1):1–10. Available from: http://dx.doi.org/10.1186/s13071-015-0987-y

15. Singh P, Kumar M. Current treatment of visceral leishmaniasis (Kala-azar): an overview. Int J Res Med Sci [Internet]. 2014;2(3):810. Available from: http://www.msjonline.org/?mno=157405

16. Okwor I, Uzonna J. Review Article Social and Economic Burden of Human Leishmaniasis. Am Soc Trop Med Hyg. 2016;94(3):489–93.

17. Collin S, Davidson R, Ritmeijer K, Keus K, Melaku Y, Kipngetich S, et al. Conflict and Kala-Azar: Determinants of Adverse Outcomes of Kala-Azar among Patients in Southern Sudan. Clin Infect Dis [Internet]. 2004;38(5):612–9. Available from: https://academic.oup.com/cid/article-lookup/doi/10.1086/381203

18. Queiroz A De, Cavalcanti NV. Risk Factors for Death in Children with Visceral Leishmaniasis. PLoS Negl Trop Dis. 2010;4(11):1–5.

19. Gerais M, Sérgio A, De AC, Juniorii CT, Rabelloiii A. Revista da Sociedade Brasileira de Medicina Tropical Factors of poor prognosis of visceral leishmaniasis among children under 12 years of age. A retrospective monocentric study in Belo. 2013;46(1):1–7.

20. Costa DL, Rocha RL, de Brito Ferreira Chaves E, de Vasconcelos Batista VG, Costa HL, Nery Costa CH. Predicting death from kala-azar: Construction, development, and validation of a score set and accompanying software. Rev Soc Bras Med Trop. 2016;49(6):728–40.

21. Belo VS, Struchiner CJ, Barbosa DS, Nascimento BWL, Horta MAP, da Silva ES, et al. Risk Factors for Adverse Prognosis and Death in American Visceral Leishmaniasis: A Meta-analysis. PLoS Negl Trop Dis. 2014;8(7):1–9.

22. Druzian AF, Souza AS d., Campos DN d., Croda J, Higa MG, Dorval MEC, et al. Risk factors for death from visceral leishmaniasis in an urban area of Brazil. PLoS Negl Trop Dis. 2015;9(8):1–11.

23. Alemayehu M., Wubshet M. MN. Magnitude of visceral leishmaniasis and poor treatment outcome among HIV patients: Meta-analysis and systematic review. HIV/AIDS - Res Palliat Care [Internet]. 2016;8:75–81. Available from: http://www.embase.com/search/results?subaction=viewrecord&from=export&id=L610074224%0Ahttp://dx.doi.org/10.2147/HIV.S96883

24. Gebreyohannes EA, Bhagvathula AS, Abegaz TM, Seid MA. Treatment outcomes of visceral leishmaniasis in Ethiopia from 2001 to 2017: A systematic review and meta-analysis. Infect Dis Poverty. 2018;7(1):1–9.

25. Assumpção Mourão MV, Toledo A, Gomes LI, Freire VV, Rabello A. Parasite load and risk factors for poor outcome among children with visceral leishmaniasis. A cohort study in Belo Horizonte, Brazil, 2010-2011. Mem Inst Oswaldo Cruz. 2014;109(2):147–53.

26. de Araújo VEM, Morais MHF, Reis IA, Rabello A, Carneiro M. Early clinical manifestations associated with death from visceral leishmaniasis. PLoS Negl Trop Dis. 2012;6(2):1–10.

27. Verma S, Kumar R, Katara GK, Singh LC, Negi NS, Ramesh V, et al. Quantification of parasite load in clinical samples of leishmaniasis patients: Il-10 level correlates with parasite load in visceral leishmaniasis. PLoS One. 2010;5(4).

28. Soares MRA, Ishikawa EAY, Silva JM, Zacarias DA, Costa DL, Costa CHN. Bone Marrow Parasite Burden among Patients with New World Kala-Azar Is Associated with Disease Severity. Am J Trop Med Hyg. 2014;90(4):621–6.

29. Das A, Karthick M, Dwivedi S, Banerjee I, Mahapatra T. Epidemiologic Correlates of Mortality among Symptomatic Visceral Leishmaniasis Cases: Findings from Situation Assessment in High Endemic Foci in India. PLoS Negl Trop Dis. 2016;DOI:10.137:1-12.

30. Desta A, Shiferaw S, Kassa A, Shimelis T, Dires S. Leishmaniasis. 2005;1–99.

31. Aderie EM, Diro E, Zachariah R, da Fonseca MS, Abongomera C, Dolamo BL, et al. Does timing of antiretroviral treatment influence treatment outcomes of visceral leishmaniasis in Northwest Ethiopia? Trans R Soc Trop Med Hyg. 2017;111(3):107–16.

32. Abongomera C, Diro E, Vogt F, Tsoumanis A, Mekonnen Z, Admassu H, et al. The Risk and Predictors of Visceral Leishmaniasis Relapse in Human Immunodeficiency Virus-Coinfected Patients in Ethiopia: A Retrospective Cohort Study. Clin Infect Dis. 2017;65(10):1703–10.

33. Croft SL, Olliaro P. Leishmaniasis chemotherapy — challenges and opportunities. Clin Microbiol Infect. 2011;17:1478–83.

34. Ministry of Health. Diagnosis and Treatment of Visceral Leishmaniasis (Kal-azar) in Kenya. 2017;2–77.

35. Sigdel B, Bhandary S, Rijal S. Epistaxis in Visceral Leishmaniasis with Hematological Correlation. Int J Otolaryngol. 2012;2012:10–3.

36. Werneck GL, Batista MSA, Gomes JRB, Costa DL, Costa CHN. Prognostic Factors for Death from Visceral Leishmaniasis in Teresina, Brazil. 2003;31(3)(3):174–5.

